# Auditory cortex activity measured with functional near-infrared spectroscopy is susceptible to masking by cortical blood stealing

**DOI:** 10.1101/2020.06.04.129205

**Authors:** Kurt Steinmetzger, Zhengzheng Shen, Helmut Riedel, André Rupp

## Abstract

To validate the use of functional near-infrared spectroscopy (fNIRS) in auditory perception experiments, combined fNIRS and electroencephalography (EEG) data were obtained from normal-hearing subjects passively listening to speech-like stimuli without linguistic content. The fNIRS oxy-haemoglobin (HbO) results were found to be inconsistent with the deoxy-haemoglobin (HbR) and EEG data, as they were dominated by pronounced cerebral blood stealing in anterior-to-posterior direction. This large-scale bilateral gradient in the HbO data masked the right-lateralised neural activity in the auditory cortex that was clearly evident in the HbR data and EEG source reconstructions. When the subjects were subsequently split into subgroups with more positive or more negative HbO responses in the right auditory cortex, the former group surprisingly showed smaller event-related potentials, less activity in frontal cortex, and increased EEG alpha power, all indicating reduced attention and vigilance. These findings thus suggest that positive HbO responses in the auditory cortex may not necessarily be a favourable result when investigating auditory perception using fNIRS. More generally, the results show that the interpretation of fNIRS HbO signals can be misleading and demonstrate the benefits of combined fNIRS-EEG analyses in resolving this issue.

## I. INTRODUCTION

Functional near-infrared spectroscopy (fNIRS) allows to derive measures of cortical activity by estimating relative concentration changes of oxy- (HbO) and deoxy-haemoglobin (HbR) in the blood. Due to its wide applicability and many practical benefits, fNIRS has become a popular neuroimaging method in recent years (Pinti et al., 2018; Scholkmann et al., 2014). At the same time, the analysis of fNIRS data has become increasingly more refined, particularly because non-cortical blood flow changes were found to strongly affect HbO measurements and methods to mitigate this confound have been introduced (Kirilina et al., 2012; Tachtsidis and Scholkmann, 2016). The aim of the current study was to validate the use of fNIRS in the context of auditory perception experiments with normal-hearing adults, by comparing fNIRS data with concurrently obtained electroencephalography (EEG) data. As previous fNIRS studies in the auditory domain mostly focussed on speech-evoked activity (Lawrence et al., 2018; Rossi et al., 2012), partly because fNIRS is especially well suited for investigating speech perception in infants (Minagawa-Kawai et al., 2011; H. Sato et al., 2012) and cochlear implant (CI) users (Anderson et al., 2017; Sevy et al., 2010), it is unclear to date if fNIRS can also be used to map more basic auditory processing. The fundamental questions in this context are whether the spatial resolution and statistical power of fNIRS are large enough to detect activity in the auditory cortex and if sufficiently strong HbO and HbR activations can be detected even when subjects do not pay attention. The latter point is of particularly relevance for possible diagnostic applications, such as the evaluation of CI-based hearing in infants.

In contrast to previous studies investigating auditory perception using combined fNIRS and EEG measurements (Chen et al., 2015; Telkemeyer et al., 2011), we compared the distributed EEG source activity to the NIRS topographies, rather than focussing on the event-related potentials only. To enable accurate source localisations, the positions of the EEG electrodes and fNIRS optodes were digitised and co-localised. Furthermore, an fNIRS optode layout was used that covered a markedly larger area of the scalp than in previous studies, enabling to evaluate the distribution and extent of the cortical activity across the entire temporal lobes and adjacent regions. To ensure that the fNIRS data were only reflecting cortical activity, we also included channels with a short source-detector separation in the montage and used them to statistically control for superficial blood flow changes in the analyses (T. Sato et al., 2016; Tachtsidis and Scholkmann, 2016).

The stimulus materials employed in the present study had speech-like acoustic properties and prosodic variations, but no linguistic content, and were intended to elicit activity in the primary auditory cortices and adjacent areas along the superior temporal lobe (Andermann et al., 2017; Gutschalk et al., 2002; Norman-Haignere et al., 2019; Patterson et al., 2002). Specifically, the sounds comprised different stimulus classes that varied with respect to their pitch strength (strong, weak, or none) and prosodic contours (static or dynamic), both of which are essential for speech comprehension and intelligibility (Oxenham, 2008; Steinmetzger and Rosen, 2015, 2018). As the prosodic information conveyed by the stimuli varied slowly over time, we furthermore expected the activity in the superior temporal lobe to be lateralised to the right (Boemio et al., 2005; Poeppel, 2003; Wartenburger et al., 2007). Due to the passive task and the relatively small cortical region expected to be activated, the design of the current experiment was intended to maximise the evoked fNIRS and EEG responses by using a block design with long stimulation periods and equally long pauses as well as a large number of trials. Additionally, the data were pooled across all stimulus conditions to maximise the data quality and to make the results generalisable.

## II. METHODS

### A. Participants

Twenty subjects (9 females, 11 males; mean age 23 years, standard deviation 2.8 years) were tested. They were all right-handed and reported no history of neurological or psychiatric illnesses. All participants used German as their main language and had audiometric thresholds of less than 20 dB hearing level (HL) at octave frequencies between 125 and 8000 Hz. All subjects gave written consent prior to the experiment and the study was approved by the local research ethics committee (Medical Faculty, University of Heidelberg).

### B. Stimuli

The experiment comprised five different stimulus conditions, which were pooled together for the analyses. The stimuli were based on recordings of 16 male talkers reading five- to six-sentence passages (Green and Rosen, 2013; Steinmetzger and Rosen, 2015, 2018). For the first condition (‘*Strong pitch – dynamic F0’*), the pitch contours of the recordings were extracted, interpolated through silent and voiceless periods, and used to generate tone complexes with dynamically varying pitch tracks in which the phases of the component tones approximated a typical adult male glottal pulse. The resulting complexes were normalised to a median fundamental frequency (*F*0) of 100 Hz. The first 12 s of each tone complex were then selected and divided into consecutive 1-second segments, resulting in a total of 192 individual stimuli (16 talkers x 12 segments). A second stimulus condition with monotonic pitch contours (‘*Strong pitch – static F0’*) was constructed by dividing the log-transformed distribution of all *F*0 values of each talker into 12 quantiles and using these values to generate another set of 192 1-second tone complexes. In the first two stimulus conditions, the frequencies of all component tones were integer multiples of the *F*0 and thus harmonically related, giving rise to a strong pitch percept. Additionally, inharmonic equivalents of the first two conditions were produced by shifting the frequencies of all component tones by 25% of the median *F*0 of each individual stimulus (‘*Weak pitch – dynamic F0*’ and ‘*Weak pitch – static F0’*). This procedure reduces the pitch strength of the stimuli (Roberts and Brunstrom, 2001), but leaves all other acoustic properties largely unchanged. The components were shifted by 25% as this value was shown to maximise the degree of inharmonicity for tone complexes with a fixed pitch (Roberts et al., 2010). For half the stimuli the shift was applied upwards, for the other half it was applied downwards. A fifth condition, in which the stimuli contained no pitch information at all (‘*No pitch’*), was based on 192 different 1-second segments of white noise.

All stimuli had a sampling rate of 48 kHz and their spectra were shaped to have the same long-term average speech spectrum (Greenwood-spaced 0.25-octave smoothing, FFT size = 512 samples), using a frequency sampling-based FIR filter (4096^th^ order, applied forwards and backwards). To further align their spectra, the stimuli were band-pass filtered between 180–4500 Hz (8^th^-order Chebyshev type 2, applied forwards and backwards, stop-band attenuation 35 dB). After applying 25-ms Hann-windowed on- and offset ramps, all stimuli were adjusted to have the same root-mean-square level.

Example stimuli of all five conditions are shown in Fig. 1A. For the waveforms depicted in the upper row, it is apparent that they are only periodic for the stimuli with a strong pitch, while they are less regular for the conditions with a weak pitch, and completely aperiodic for the stimuli with no pitch. The narrow-band spectrograms shown in the middle row demonstrate that the spectra of the stimuli are indeed very similar, despite the markedly different waveforms. The lower row shows spectrographic representations of summary autocorrelation functions (SACFs; Meddis and Hewitt, 1991; Meddis and O’Mard, 1997), for which individual autocorrelation functions were calculated for the low-pass filtered (2^nd^-order Butterworth, cut-off 1 kHz) outputs of 22 gammatone filters with centre frequencies ranging from 0.2–4 kHz and summed together into SACFs. This procedure was applied with a step size of 1 ms and a Hann-window size of 5 ms to yield spectrographic representations of the SACFs across the duration of the stimuli. Importantly, while the lag time of the first peak in the SACF spectrograms represents the *F*0 of the stimuli, the height of this peak may be interpreted as a measure of pitch strength (Yost et al., 1996). In line with this notion, the peak around 10 ms is visibly more pronounced and symmetric for the stimuli with a strong pitch compared to those with a weak pitch. Due to the absence of any temporal regularity, no such peak was present for the stimuli with no pitch.

**Figure 1.**
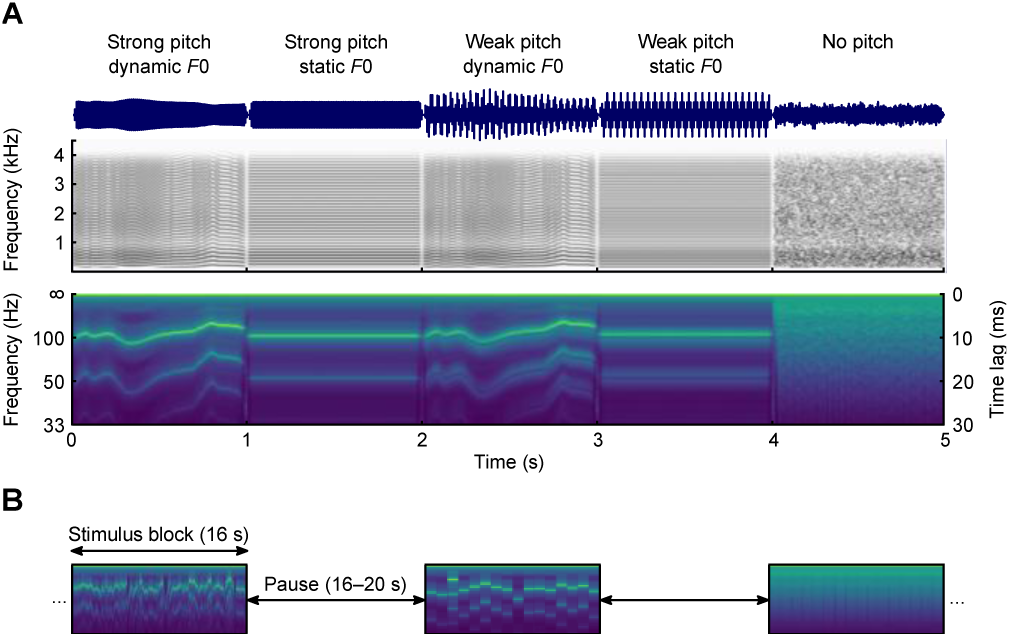
Example stimuli and experimental design. A) Waveforms, narrow-band spectrograms, and summary autocorrelation function spectrograms of example stimuli of the five conditions. B) The individual 1-s stimuli were presented in blocks of 16, alternating with pauses lasting 16–20 s.

### C. Procedure

As haemoglobin concentration changes evolve over the course of several seconds, a block design with long stimulation periods and equally long pauses was used to maximise the experimental effects. The individual stimuli within each condition were thus randomly concatenated into blocks consisting of 16 stimuli. The stimulus blocks were followed by pauses with random durations ranging from 16–20 s. Each participant was presented with 12 blocks in each of the 5 experimental conditions, adding up to 192 individual stimuli per condition. The order of the blocks was randomised without any constraints. The experiment consisted of 60 stimulus blocks framed by 61 pauses, amounting to a total duration of about 34 mins. As shown by the SACF spectrograms in Fig. 1B, stimulus blocks with a dynamic *F*0 had continuously changing pitch contours, while those with a static *F*0 had step-like pitch patterns.

The experiment took place in a sound-attenuating and electrically shielded room, with the participant sitting in a comfortable reclining chair during data acquisition. Throughout the experiment, the light was dimmed to the lowest level to minimise ambient light from interfering with the NIRS recordings. There was no behavioural task, but pauses were inserted about every 10 mins to ensure the vigilance of the subjects. The stimuli were presented with 24-bit resolution at a sampling rate of 48 kHz using an RME ADI-8 DS sound card (Haimhausen, Germany) and presented via Etymotic Research ER2 earphones (Elk Grove Village, IL, USA) attached to a Tucker-Davis Technologies HB7 headphone buffer (Alachua, FL, USA). The presentation level was set to 70 dB SPL, using an artificial ear (Brüel & Kjær, type 4157, Nærum, Denmark) connected to a corresponding measurement amplifier (Brüel & Kjær, type 2610, Nærum, Denmark).

### D. fNIRS recording and analysis

fNIRS signals were recorded with a continuous-wave NIRScout 16×16 system (NIRx Medizintechnik, Berlin, Germany) at a sampling rate of 7.8125 Hz. Eight source optodes and eight detector optodes were placed symmetrically over each hemisphere by mounting them on an EEG cap (EasyCap, Herrsching, Germany). The source optodes emitted infrared light with wavelengths of 760 and 850 nm. To avoid interference between adjacent sources, only a single source optode per hemisphere was illuminated at a given time. The chosen optode layout was devised to optimally cover the auditory cortex and associated areas, resulting in 22 measurement channels per hemisphere, of which 20 had a standard source-to-detector distance of about 30 mm, while the remaining 2 had a shorter spacing of about 15 mm. The optode and reference positions for each individual subject were digitised with a Polhemus 3SPACE ISOTRAK II system (Colchester, VT, USA) before the experiment.

The data were pre-processed using the HOMER2 toolbox (version 2.8; Huppert et al., 2009) and custom MATLAB (MathWorks, Natick, MA, USA) code. The raw light intensity signals were first converted to optical density values and then corrected for motion artefacts. A kurtosis-based wavelet algorithm with a threshold value of 3.3 (Chiarelli et al., 2015) was used to identify and correct motion artefacts by rejecting spectral components of the signal rather than time segments. Measurement channels with poor signal quality were then excluded from further analysis based on their scalp coupling index (SCI; Pollonini et al., 2014). SCIs were computed by filtering the optical density signals of both wavelengths between 0.5–2.5 Hz (3^rd^-order low-pass and 5^th^-order high-pass Butterworth filters, applied forwards and backwards), to emphasise the heart-beat related signal fluctuations, and correlating the filtered signals. Channels with correlation coefficients below 0.75 were excluded (mean = 1.25 channels/subject, max. 6 per subject), as this indicates a poor contact between optodes and scalp. Next, the motion-corrected signals of the remaining channels were band-pass filtered between 0.01–0.5 Hz (same filter types as for the SCI above), to isolate the task-related neural activity, and subsequently converted to concentration values based on the modified Beer-Lambert law (Scholkmann et al., 2014). The differential path length factors required for the conversion were determined based on the wavelengths of the light and the age of the subject (Scholkmann and Wolf, 2013).

Secondly, the pre-processed data were statistically evaluated and topographically visualised with SPM-fNIRS (version r3; Tak et al., 2016). Based on the principles of the general linear model (GLM), the SPM framework tests how closely the stimulus-evoked signal changes resemble a canonical hemodynamic response function (HRF). In SPM-fNIRS, the optode positions of each subject were first transformed from subject space to MNI space, after which they were probabilistically rendered onto the ICBM-152 cortical template surface. This was achieved by applying a least-squares approach in which a set of digitised reference points distributed across the whole head (4 external fiducials and 17 positions of the extended international 10-20 system) was aligned to template values (Singh et al., 2005). The signals were then temporally smoothed using the shape of the canonical HRF waveform (‘pre-colouring’) to avoid autocorrelation issues when estimating the model (Worsley and Friston, 1995). The data of the individual subjects were statistically modelled by convolving the continuous signals obtained from each long channel with separate regressor functions for each of the five experimental conditions. The standard SPM double-gamma function was used as canonical HRF and convolved with 16-s boxcar functions following the onsets of the stimulus blocks. The HbO data were modelled with positive HRFs, while the concentration changes were assumed to be negative for the HbR analysis. To allow the time course of the measured concentration changes to vary slightly, the temporal and spatial derivatives of the canonical HRF were included as additional regressors (Plichta et al., 2007). Furthermore, the first component of a principal component analysis of the pre-processed signals of the four short channels was used as an additional nuisance regressor, as this serves to estimate and remove the so-called global scalp-hemodynamic component (T. Sato et al., 2016), i.e. the superficial signal component. After estimating the HbO and HbR GLMs for each subject, a contrast vector was defined in which the regressors for the five experimental conditions were set to 1, whereas the regressors representing the derivatives and the global scalp component were set to 0 to statistically control for their effects. Group-level statistics were then computed for each long channel by testing whether the subject-level β weights within a given group were significantly greater than 0. Non-parametric right-tailed Wilcoxon signed-rank tests were used throughout, as Kolmogorov-Smirnov tests indicated that the β weights for the HbO and HbR data had non-normal distributions for most long channels. If two subject groups were compared, non-parametric right-tailed Wilcoxon rank-sum tests were employed.

A customised version of the SPM-fNIRS plotting routine was devised to topographically visualise the optode and channel positions as well as the functional activations. The MNI-transformed optode and channel locations as well as the measured channel lengths are shown in Fig. 2. The short channels were omitted in the plots of the functional activations (Figs. 3 & 4) as they are assumed to not reflect any cortical responses. In any plots depicting HRFs (Figs. 3, 4 & 6), the waveforms were averaged from -2–32 s around block onset and baseline corrected by subtracting the mean amplitude in the pre-stimulus window from each sample point. The corresponding HRFs are shown both after the pre-processing (‘Total HRF’) and after regressing out the contribution of the short channels and pre-colouring the signals (‘Cortical HRF’), to illustrate the effect of removing the non-cortical signal component. Additionally, the β weighted canonical HRFs (‘Model HRF’) are shown to demonstrate how well the cortical HRFs can be explained by the HRF models.

**Figure 2.**
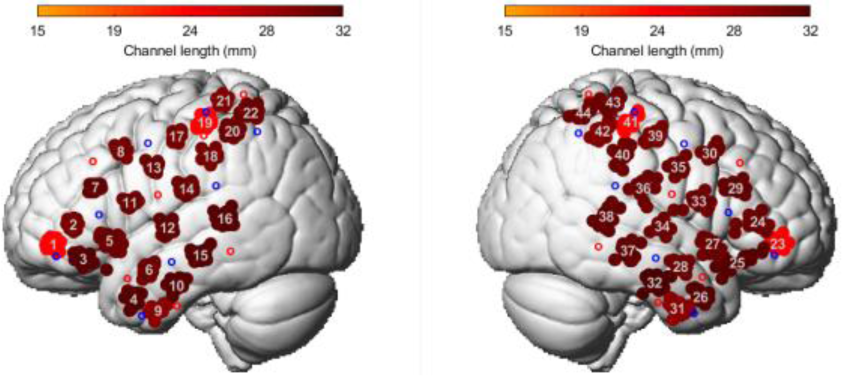
fNIRS optode and channel layout. The average MNI positions of the source and detector optodes are shown as red and blue circles, respectively. The average positions of the resulting measurement channels are indicated by the grey numbers. The point clouds around each channel signify the MNI positions for the individual subjects and the colour of the clouds shows the average channel length, i.e. the mean distance between the corresponding source-detector pair.

**Figure 3.**
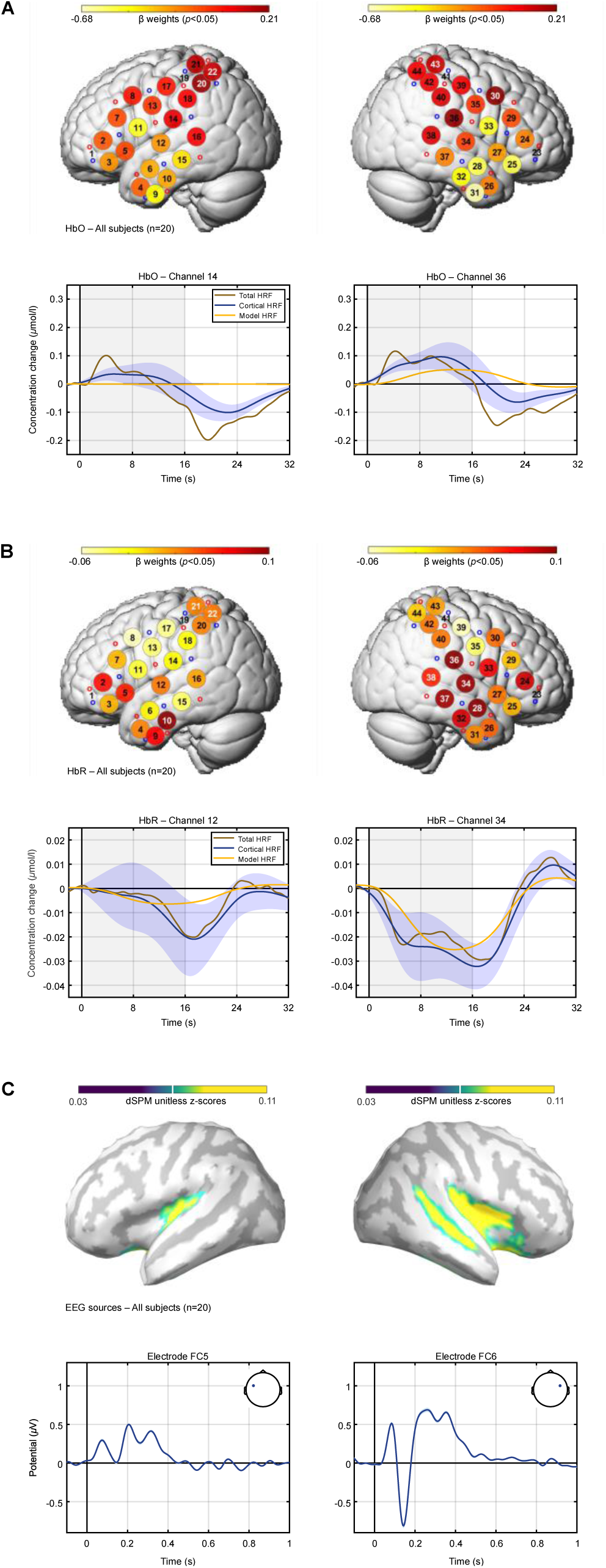
Group results. A) fNIRS HbO results averaged across all 20 participants. The upper row shows the topographical activation patterns and the lower row the hemodynamic response functions (HRFs) for two channels covering the left and right auditory cortex. White channel numbers in the topographies indicate significance. B) fNIRS HbR results. C) ERPs recorded from two electrodes over the left and right hemispheres and dSPM EEG source reconstructions across the whole stimulus duration. In the EEG source plots, only cortical surface points with at least 50% of the maximal amplitude are shown in colour. The shading around the fNIRS and EEG time courses indicates the standard error of the mean. A strong lateralisation to the right hemisphere was observed in all three analyses.

**Figure 4.**
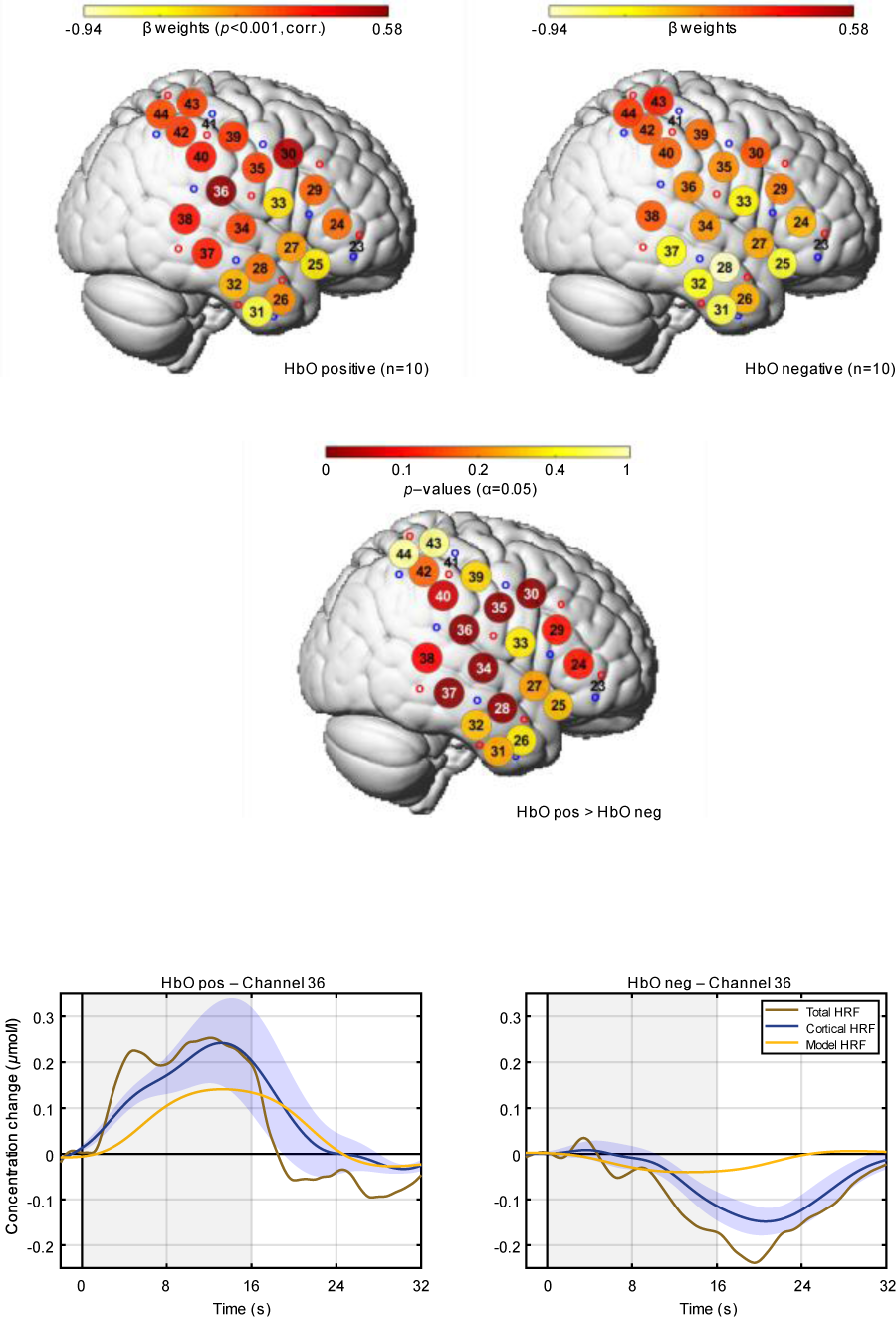
HbO subgroups: fNIRS results. fNIRS HbO topographies and HRFs after dividing the subjects into subgroups with more positive or more negative HbO responses in the right auditory cortex. Stronger activity in the right hemisphere was observed for the HbO positive group.

**Figure 5.**
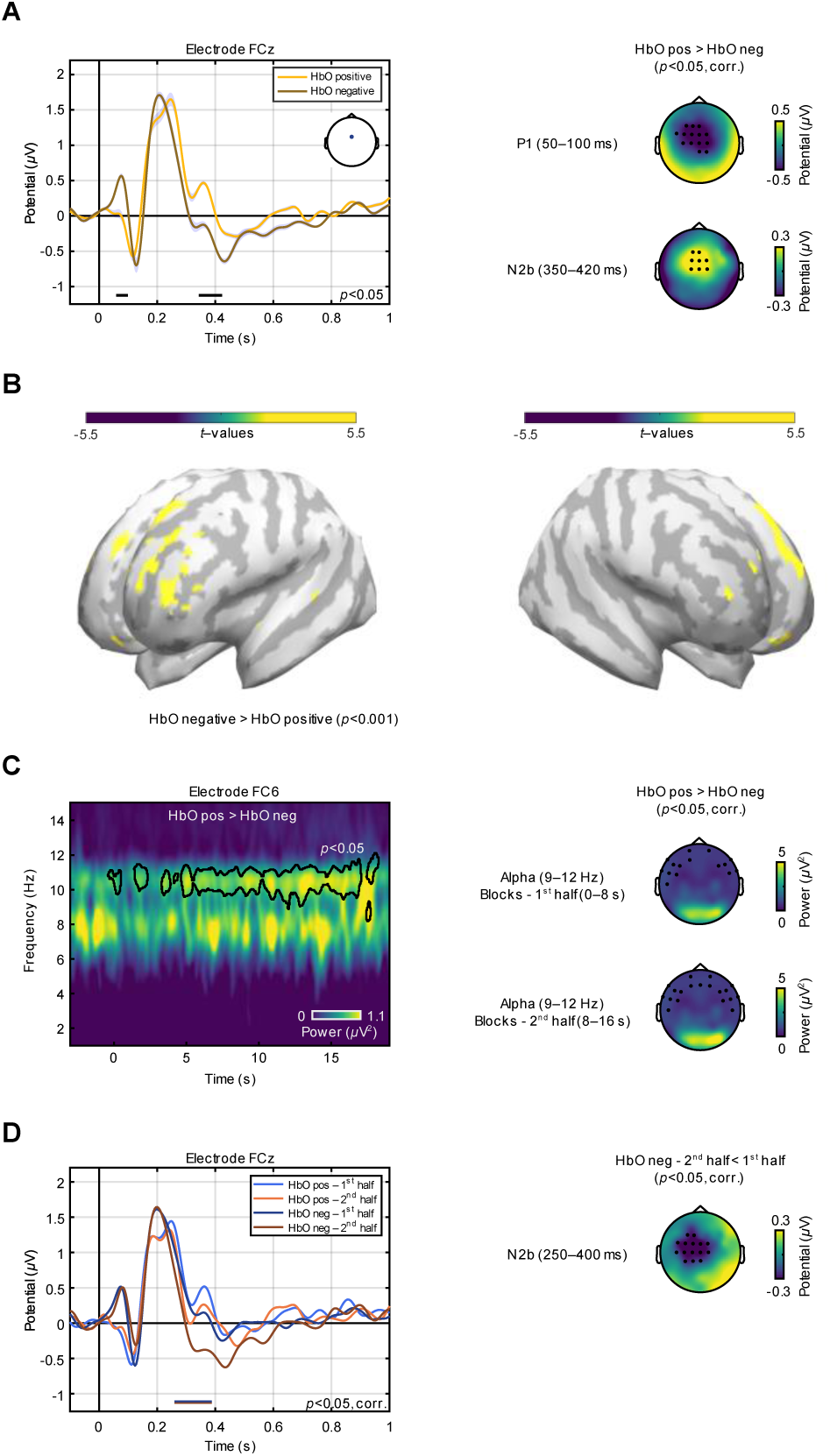
HbO subgroups: EEG results. A) ERPs of the HbO subgroups. Amplitudes were smaller in the HbO positive group, as indicated by the ERP traces (left) and the corresponding scalp maps (right). Significant time windows are indicated by the horizontal black bars below the ERPs. The scalp maps on the right show group differences of the time-averaged voltages. Electrodes for which the difference between groups was significant are indicated by black dots. B) dSPM EEG source reconstructions. Stronger activity in the left prefrontal cortex was observed in the HbO negative group. C) Absolute EEG power in the upper alpha band increased throughout the stimulus blocks for the HbO positive group. On the left, a time-frequency plot shows the difference between the subgroups, where the area outlined by the black line indicates significance. The scalp maps on the right demonstrate that this effect was significant at frontal and temporal sites and became stronger in the second half of the blocks for electrodes over the right hemisphere. D) ERPs in the first and second halves of the stimulus blocks. A significant difference between the first and second halves was observed in the HbO negative group, as indicated by the horizontal bar. The scalp map on the right shows the topography of this difference.

**Figure 6.**
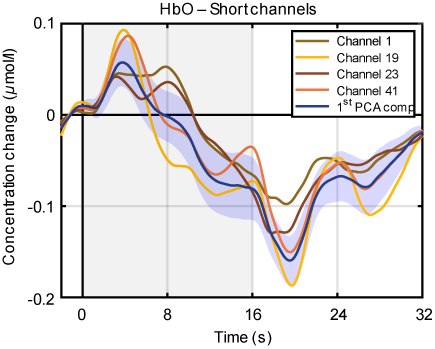
HbO signals of the short channels. The HRFs of the four short channels are shown together with their first PCA component. The PCA amplitude was reduced by a factor of 4 so that all signals exhibit comparable amplitudes. Note that all four short channels have very similar time courses, despite their large spatial separation.

### E. EEG recording and analysis

Continuous EEG signals were recorded using a BrainVision actiCHamp system (Brain Products, Gilching, Germany) with 60 electrodes arranged according to the extended international 10-20 system. Four additional electrodes were placed around the eyes to record vertical and horizontal eye movements. The EEG data were recorded with an initial sampling rate of 500 Hz, an online anti-aliasing low-pass filter with a cut-off frequency of 140 Hz and were referenced to the right mastoid. As with the fNIRS optodes, the electrode positions of each subject were digitized with a Polhemus 3SPACE ISOTRAK II system before the experiment.

The data were pre-processed offline using FieldTrip (version 20180924; Oostenveld et al., 2011) and custom MATLAB code. The continuous waveforms were first segmented into epochs ranging from -0.3–1.1 s around stimulus onset. Next, the epochs were re-referenced to the mean of both mastoids and detrended as well as demeaned by removing a 1^st^-order polynomial. The epochs were then low-pass filtered (cut-off 15 Hz, 4^th^-order Butterworth, applied forwards and backwards), baseline corrected by subtracting the mean amplitude from -0.1–0 s before stimulus onset, and subsequently down-sampled to 250 Hz. After visually identifying and excluding bad channels (total = 4, max. 2 per subject), the data were decomposed into 20 principal components to detect and eliminate eye artefacts. After the 4 eye electrodes were removed from the data, epochs in which the amplitudes between -0.2–1 s around stimulus onset exceeded ±60 µV or the z-transformed amplitudes differed by more than 15 standard deviations from the mean of all channels were excluded from further processing. On average, 86% of the trials (830/960 per subject, min. 65% per subject) passed the rejection procedure. Lastly, bad channels were interpolated using the weighted average of the neighbouring channels and the data were re-referenced to the average of all 60 channels.

Grand-average event-related potentials (ERPs) were computed by applying a baseline correction from -0.1–0 s before stimulus onset to the individual trials, averaging the epochs of each subject, and calculating the group mean of the subject-level ERPs. The ERPs were statistically examined using non-parametric permutation tests (Maris and Oostenveld, 2007). To compare the ERPs between two groups, independent- or dependent-samples *t*-tests were calculated for all combinations of electrodes and sample points during the stimulation period (0–1 s). The scalp distributions of the ERP components, in contrast, were tested by first averaging the voltages over time. Where possible, false positive findings were controlled for by merging significant samples into temporally and spatially coherent clusters, if at least 3 neighbouring electrodes showed the respective effect. The *t*-values in each cluster were then summed to derive a test statistic, which was compared to the same statistic after the trials were randomly re-allocated to the two groups. This step was repeated 5000 times to obtain the *p*-value of a cluster. Uncorrected tests were computed in the same way, but the individual *t*-values were re-allocated between groups to derive their *p*-values.

Distributed source reconstructions of the averaged ERPs were computed using the MNE-dSPM approach implemented in Brainstorm (version 29-Apr-2020; Dale et al., 2000; Tadel et al., 2011). The electrode positions of each subject were co-registered to the ICBM152 MRI template by first aligning three external fiducial points (LPA, RPA, and Nz) and subsequently projecting the electrodes to the scalp of the template MRI. A Boundary Element Method (BEM) volume conduction model based on the ICBM152 template and the corresponding cortical surface (down-sampled to 15.000 vertices) were used as head and source models. The BEM head model was computed using OpenMEEG (version 2.4.1; Gramfort et al., 2010) and comprised three layers (scalp, outer skull, and inner skull) with 1082, 642, and 642 vertices, respectively. Linear MNE-dSPM solutions with dipole orientations constrained to be normal to the cortex were estimated after pre-whitening the forward model with the averaged noise covariance matrix calculated from the individual trials in a time window from -0.2–0 s before stimulus onset. The default parameter settings for the depth weighting (order = 0.5, max. amount = 10), noise covariance regularisation (regularise noise covariance = 0.1), and regularisation parameter (SNR = 3) were used throughout. The individual source reconstructions were then converted to absolute values, spatially smoothed (Gaussian kernel, full width at half maximum = 3 mm), and averaged across subjects and time points. Statistical differences between groups were evaluated using non-parametric permutation tests, as described above.

For the time-frequency analysis, ERPs across the duration of the blocks were computed by extracting epochs ranging from -4.1–20.1 s around the onset of the first stimulus in each block. Pre-processing steps that did not differ from the procedure used for the individual stimuli are omitted for brevity. Here, a high-pass filter (cut-off 1 Hz, 3^rd^-order Butterworth, applied forwards and backwards) was used to eliminate slow drifts and epochs with amplitudes exceeding ±100 µV between -4–20 s or z-transformed amplitudes that differed by more than 30 standard deviations from the mean were rejected. On average, 93% of the trials (56/60, min. 70% per subject) passed the rejection procedure.

The pre-processed individual stimulus blocks were time-frequency transformed by convolving the data with tapered wavelets in the frequency domain. A Hanning-windowed taper with a constant length of 1 s was used across all frequencies. The frequency range of the decomposition was set to 1–15 Hz, with a resolution of 0.05 Hz, and the time window ranged from -4–20 s, with a step size of 0.05 s. The length of the blocks was zero-padded to the next power of 2 for computational efficiency. The resulting time-frequency representations were then averaged across blocks to derive estimates of the total EEG power, i.e. evoked and induced activity, for each subject. No baseline correction was applied to the total EEG power estimates to retain possible differences between subject groups in the pre-stimulus period. Statistical differences between groups were either assessed across electrodes, frequencies, and time points, or after averaging across frequency and time windows to enable a topographic representation of the results. In both cases, the same non-parametric permutation tests as described above were used.

## III. RESULTS

### A. Group results

The grand-averaged fNIRS HbO data of all 20 subjects surprisingly showed no significant activation of the auditory cortices, despite a trend in the right hemisphere (Fig. 3A). The clearest sign of an activation was observed at channel 36 (*p* = 0.16), which covered the posterior part of the right auditory cortex. However, the HRF at this channel peaked markedly earlier and receded before the canonical HRF model, implying that other processes in addition to a standard blood-oxygen-level-dependent (BOLD) response were involved. Instead of the expected auditory cortex activation, the HbO data exhibited a bilateral anterior-to-posterior gradient, with decreased activity in frontal areas but increasingly more positive responses towards posterior regions. A simple linear regression model that included the positions of the long NIRS channels as sole predictor, confirmed that the β weights were higher the more posterior a channel was positioned (*t* (776) = 4.5, *p* < 0.001). In line with this finding, several channels in this cortical region displayed significant activity (*p* < 0.05; chs. 20, 22, 30 & 43).

The fNIRS HbR data, in contrast, showed the expected activation of the auditory cortex, albeit strongly lateralised to the right hemisphere (Fig. 3B). As is typical for HbR measurements, the changes in haemoglobin concentration were markedly smaller compared to the HbO data, which is also reflected in the reduced β weights of the channels. Nevertheless, a cluster of five channels distributed across the right auditory cortex showed significant activity (*p* < 0.05; 28, 34 & 36–38), with the strongest effect observed at channel 34 (*p* = 0.006), located near the primary auditory cortex. Furthermore, the cortical HRF for channel 34, closely matched the shape of the model HRF. In agreement with prior findings (Kirilina et al., 2012), the contribution of the superficial signal component was also markedly smaller than for the HbO data, as indicated by the greater similarity of the total and cortical HRFs.

In line with these findings, a strong lateralisation to the right was also evident in the EEG data, both at the sensor and source level (Fig. 3C). The ERPs recorded over the left auditory cortex were much smaller compared to those recorded over the right one. Moreover, the distributed source reconstructions of the activity evoked across the entire duration of the individual stimuli also exhibited little activity in the left auditory cortex, along with a focus of activity around the right auditory cortex and superior temporal sulcus.

Taken together, all three analyses coincide regarding the strong lateralisation of the neural activity to the right, with negligible responses in the left hemisphere. The HbO results, however, deviate from the HbR and EEG data in that a pronounced activation of auditory cortex is seemingly lacking altogether.

### B. HbO subgroup results

The evaluation of the fNIRS HbO data on the subject level showed strongly increased activity in the right auditory cortex for some participants, but also pronounced deactivations for several others. Based on this observation, a median split was performed to assign the subjects into equally sized subgroups with more positive or more negative HbO responses, respectively. To achieve this, the baseline corrected cortical HRFs of the channels closest to the right primary auditory cortex (nos. 34 & 36) were averaged for each subject and the area under the curve across the duration of the stimulation period (0–16 s) was calculated, i.e. the period during which the HRFs was expected to rise and peak.

The fNIRS topographies in each subgroup and their statistical comparison (Fig. 4) demonstrate that the 10 subjects in the HbO positive group indeed showed a much greater activation of the right auditory cortex, with a maximum at channel 36 in both cases (*p*<0.05, Holm-Bonferroni corrected). With a lowered significance threshold (*p*<0.05, uncorrected), the subgroup comparison resulted in differences across the right auditory cortex, in line with the HbR group data, as well as additional activity beyond the right auditory cortex (chs. 30, 35 & 40) and in the left hemisphere (chs. 3 & 16, not shown). In contrast, no differences between the subgroups were observed at this significance level in the HbR data. Crucially, whereas the cortical HRF of the HbO positive group for channel 36 closely resembled the shape of the HRF model, the cortical response in the HbO negative group deviated markedly from the model function. Here, the negative peak of the cortical HRF occurred several seconds after stimulus offset and hence much later than expected. In summary, it appears that the positive and negative responses in the two subgroups cancelled each other out on group level, explaining the unexpectedly small HbO activations.

Next, it was evaluated whether the ERPs also differed across the two HbO subgroups (Fig. 5A). Contrary to what the NIRS data might have suggested, both the P1 and the N2 were largely absent in this subgroup, while both components were present in the HbO negative group. A statistical comparison returned two significant time windows (*p*<0.05), the first one in the P1 range (∼50– 100 ms) and the second one during the N2 window (∼350–420 ms), during which the ERP amplitudes were smaller in the HbO positive group. In the corresponding scalp maps, the mean amplitudes in the negative HRF group have been subtracted from those in the positive HRF group to illustrate the topographies of these differences. For both time windows, the group difference was most pronounced in the fronto-central scalp region, as is typical for auditory ERPs. Statistical tests comparing the time-averaged amplitudes in the two groups returned a significant cluster of electrodes (*p*<0.05, cluster-corrected) in this region for both time windows.

The comparison of the EEG source reconstructions of the two subgroups (Fig. 5B) showed the strongest differences in prefrontal cortex (*p*<0.001) rather than auditory areas. The left prefrontal cortex in particular exhibited stronger activity in the HbO negative subgroup. The region at the centre of the significant area appears to be the left dorsolateral prefrontal cortex, which has been associated with top-down attentional control (Fletcher et al., 1998; Hopfinger et al., 2000; MacDonald et al., 2000).

Since the EEG sources indicated a possible difference in attention between the two subgroups, it was also examined whether the EEG alpha power differed across groups. Indeed, the absolute power of the EEG signals in the upper alpha band (∼9–12 Hz) was found to be higher in the HbO positive group. This effect increased over the course of the stimulus blocks over right temporal sites, such as electrode FC6 (Fig. 5C), suggesting a progressive disengagement of the affected regions caused by attentional top-down effects (Jensen and Mazaheri, 2010; Strauß et al., 2014). At this location, a significant group difference (*p*<0.05) was observed throughout the second half of the blocks but occurred only sporadically during the initial half. Hardly any significant differences were observed before or after the stimulus blocks as well as outside the upper alpha band, despite an evident trend in the lower alpha region. This effect was topographically evaluated by separately averaging the data in the first and second halves of the blocks across the upper alpha band. A statistical comparison of the groups (*p*<0.05, cluster-corrected) indicated higher alpha power in the HbO positive group for a large cluster of electrodes in the frontal and temporal scalp region during both halves. However, an increase in alpha power from the first to the second half was only observed over the right hemisphere, in line with the strong lateralisation found in both the fNIRS and EEG data.

Based on this finding, it was also evaluated whether the ERPs in the two groups differed between the first and second halves of the blocks. Here, the ERPs of the individual stimuli were sorted according to which half they appeared in, omitting the first stimulus in each block because of its large energy onset response. The resulting ERPs (Fig. 5D) showed no significant differences, apart from an effect during the N2 window for the HbO negative group (∼250–400 ms), where ERP amplitudes were larger during the second half (*p*<0.05, cluster-corrected). The scalp distribution of this effect shows that a cluster of electrodes with a central, slightly left-lateralised location exhibited a significant difference throughout this time window (*p*<0.05, cluster-corrected). Given its distribution and latency, this effect appears to be an N2b component that is usually evoked if listeners voluntarily pay attention to the stimuli and detect a deviation from the preceding auditory context (Patel and Azzam, 2005).

## IV. DISCUSSION

To validate the usefulness of fNIRS in the context of auditory perception studies, combined fNIRS and EEG data were obtained from normal-hearing listeners. In summary, the passive auditory paradigm employed in the current study was found to elicit consistent right lateralised fNIRS HbR and EEG responses. The fNIRS HbO data, in contrast, deviated markedly from this pattern and revealed no significant activation of the right auditory cortex on group level. Instead, the HbO results showed a bilateral anterior-to-posterior gradient, with pronounced negative responses in frontal and temporal areas, accompanied by positive responses in more posterior areas.

However, while some subjects showed negative responses in the right auditory cortex, others were found to exhibit positive HbO responses in this region. To further investigate this difference, the subjects were split into evenly sized subgroups, based on the their HbO responses in the right auditory cortex. Subsequent analyses returned the seemingly puzzling finding that larger HbO responses went along with smaller ERP amplitudes, less activity in frontal cortex, and increased EEG alpha power. These effects all suggest reduced attention in the HbO positive subgroup.

In line with the literature on the physiological bases of fNIRS signals (e.g. Kirilina et al., 2012), the HbR responses were found to be affected by extracerebral sources to a lesser extent, but they were also much smaller in amplitude than the HbO data. Hence, despite the very good fit of the measured cortical HRFs and the canonical HRF model, their statistical power was limited, as indicated by smaller *p*-values in comparison to the HbO subgroup analyses. This underlines the need to further investigate the factors that were involved in generating the unexpected HbO results.

### A. The anterior-to-posterior gradient in the HbO data is due to ‘blood stealing’

Negative BOLD responses have repeatedly been observed on a local scale and are taken to reflect the inhibition of irrelevant neural populations. Such deactivations have, for example, been found in parts of the occipital cortex in response to visual stimuli (Maggioni et al., 2015; Shmuel et al., 2002), the ipsilateral sensorimotor cortex after somatosensory stimulation (Mullinger et al., 2014), and the primary auditory or visual cortices when attention was guided to the other modality (Shomstein and Yantis, 2004). Mullinger et al. (2014) also found that negative BOLD responses in the sensorimotor cortex were accompanied by greater alpha power and larger ERPs in this region. Yet, although the ERPs were larger in the negative HbO subgroup, in line with these results, alpha power was not increased in this group, arguing against an inhibition process.

Negative BOLD effects were also observed on a larger spatial scale. Visual imagery, for instance, was reported to result in negative BOLD responses across the auditory cortex (Amedi et al., 2005) and visual stimulation was shown to cause negative BOLD effects across the auditory cortex and the default mode network (Mayhew et al., 2013), the latter of which is known to be deactivated by a wide variety of tasks (Raichle, 2015). Similarly, negative BOLD responses have been observed in the ventral fronto-parietal attention system in the right hemisphere and are thought to reflect sustained attention (Corbetta et al., 2008). Yet, while the gradient in the HbO data clearly is not a local effect, it also does not coincide with the distribution of these two networks. Although negative HbO responses where observed in frontal areas, no similar decrease was observed in the parietal areas belonging to both networks. Furthermore, although visual imagery effects cannot be ruled out entirely due to the passive task, the EEG source localisations indicated no pronounced activity in occipital areas.

Alternatively, the anterior-to-posterior gradient may be due to so-called ‘blood stealing’ or ‘vascular stealing’, which is a decrease in absolute blood flow and volume without any underlying change in neuronal activity (Harel et al., 2002; Shmuel et al., 2002), although the contribution of this process to local negative BOLD responses has repeatedly been claimed to be limited in fMRI studies (Mullinger et al., 2014; Smith et al., 2004). For the large-scale gradient in the current study, in contrast, blood stealing represents a convincing explanation because task-evoked decreases in cerebral blood volume and flow have been found to lag behind negative cortical BOLD responses (Harel et al., 2002; Shomstein and Yantis, 2004). In particular, the cortical HRFs showing negative HbO responses consistently reached their peak distinctly later then the canonical HRF model (Fig. 4). This provides strong evidence for the assumption that the negative HbO responses observed in the current study were not driven by a decrease in neural activity.

Furthermore, the HRFs of the four short channels, which reflect the non-cortical superficial signal component of the HbO data, have very similar time courses despite their large spatial separation (Fig. 6). This argues in favour of the PCA-based method used to regress out this signal component (T. Sato et al., 2016), which relies on the assumption that the superficial signals are consistent across the scalp. Moreover, this also shows that a greater number of short channels in the current fNIRS layout would not have led to a more accurate estimation of the cortical HbO signals. The most crucial feature of the short channel signals is, however, that they bear almost no resemblance with the cortical HRFs extracted from the long channels. This makes it very unlikely that the anterior-to-posterior gradient is a superficial effect and instead suggests that it is a result of cortical ‘blood stealing’.

### B. The EEG data suggest less attention in the subgroup with positive HbO responses

Three separate effects in the EEG data suggested that the different HbO results in the two subgroups are due to reduced attention in the HbO positive group: Firstly, the EEG source localisations (Fig. 5B) indicated that the left dorsolateral prefrontal cortex showed stronger activity in the HbO negative group, a region associated with top-down attentional control. For example, instructions for a subsequent task result in increased activity of this region (Hopfinger et al., 2000; MacDonald et al., 2000), while distraction from a task attenuates activity in this area (Fletcher et al., 1998). As the subjects in the current study were instructed to listen to the stimuli attentively, greater activity in this region may thus be taken to reflect increased attention.

Secondly, alpha power was higher at frontal and temporal electrode sites in the HbO positive group, and this effect increased over the right hemisphere throughout the course of the stimulus blocks (Fig. 5C), which coincided with the right-lateralisation also observed in the ERPs and EEG source localisations. Task-related alpha power increases are thought to reflect active inhibition of sensory processing, caused by attentional top-down effects (Jensen and Mazaheri, 2010; Mazaheri et al., 2014; Strauß et al., 2014). The increased alpha power in this group may thus reflect the attempt to suppress stimulus processing.

Finally, the comparison of the ERPs in the first and second halves of the stimulus blocks revealed a prominent negative deflection around 250–400 ms post stimulus onset in the HbO negative group during the second half of the blocks (Fig. 5D). Latency, distribution, and the cognitive processes involved suggest that this deflection represents an N2b component, which is typically evoked by deviating stimuli if voluntary attention is paid to the preceding auditory context (Näätänen and Gaillard, 1983; Patel and Azzam, 2005; Sussman et al., 2003). Given that the blocks with a strong or weak pitch, which made up 80% of the stimulus material, were formed from individual stimuli drawn at random from a distribution varying in terms of the pitch contours and *F*0s, some of the stimuli will inevitably have qualified as outliers based on their pitch properties. Hence, an N2b was only observed during the second half of the blocks, after the acoustic context had been sufficiently established to enable violations of expectancy. The absence of an N2b component in the HbO positive group, on the other hand, could be explained by their reduced attention to the stimuli. A similar effect has recently been reported in an magnetoencephalography (MEG) study by Fan et al. (2017), where the auditory evoked fields of Chinese listeners in response to a meaningful /ma/ sound showed a much more pronounced deflection during this latency window compared to German subjects, whose attention likely was not drawn by this stimulus to a similar extent. Interestingly, for the German group as well as the HbO positive group in the current study, the previous P2 exhibited a prominent double peak.

Despite the evidence for an attention effect presented in this section, it remains unclear how lower attention can explain the positive HbO concentration changes observed in the HbO positive group. One possibility is that the influx of oxygenated blood far exceeded the oxygen consumption in the affected areas, which together led to a strong increase in HbO concentration (Mullinger et al., 2014). Although the influx of oxygenated blood is generally thought to over-compensate the consumption of oxygen resulting from neural activity (e.g. Scholkmann et al., 2014), it is conceivable that the suppression of neural activity observed in the HbO positive group indeed caused a decrease in oxygen consumption, while the inflow of oxygenated blood continued.

### C. Absence of the P1 in the HbO positive group may signal reduced activation levels

A further notable difference between the two HbO subgroups was the absence of a discernible P1 component in the ERPs of the HbO positive group (Figs. 5A & D). Several studies have reported a link between P1 amplitude and levels of arousal and sensory activation. For example, the auditory P1 disappears during slow wave sleep (Erwin and Buchwald, 1986). Similarly, the P1 was found to be absent when subjects were injected with substances inhibiting the ascending reticular activation system (Buchwald et al., 1991). This intervention rendered the participants drowsy but awake, a state that the passive stimulation paradigm in the current study may also have induced in the subjects belonging to the HbO positive subgroup. Ultimately, the absence of the P1 thus points in the same direction as the other subgroup differences in the EEG data, namely a reduced level of attention and vigilance.

### D. Right-lateralisation of fNIRS and EEG data is due to slowly changing prosodic features

In general, the present results agree well with previous studies that have found right-lateralised responses to tones and melodies using fMRI (Norman-Haignere et al., 2019; Patterson et al., 2002) and right-lateralised EEG source localisations in response to speech sounds (Dimitrijevic et al., 2013), although the lateralisation in the present study was particularly pronounced. However, this finding is consistent with the asymmetric sampling hypothesis (Poeppel, 2003), which explicitly postulates that the slowly varying prosodic features of speech sounds should lead to strongly right-lateralized processing. This theoretical model further assumes that responses in the primary auditory cortex are largely symmetric and only secondary areas in the superior temporal cortex exhibit a strong lateralisation to the right when slow spectro-temporal changes in the stimuli are processed. This view is supported by the current fNIRS and EEG results that both showed broadly distributed activity in this region.

### E. The limited depth sensitivity of the fNIRS data did not bias the results

The light emitted by the fNIRS source optodes is believed to reach as deep into the head as about half the source-detector distance (Patil et al., 2011). For a the standard channel length of 30 mm, the scalp-brain distance in auditory areas has been shown to be mostly within the assumed 15-mm limit (Cui et al., 2011; Okamoto et al., 2004). However, MEG dipole analyses have usually indicated that the sources of electrical activity in the primary auditory cortex were somewhat retracted from the immediate cortical surface (e.g. Gutschalk et al., 2002). For example, in a combined MEG and fNIRS study, Ohnishi et al. (1997) reported an MEG dipole in the left auditory cortex that was 25 mm away from scalp surface when a 1-kHz tone was presented. It is hence very likely that parts of the activity emanating from the primary auditory cortices were not detected using the current fNIRS layout. However, since the EEG source localisations generally agree very well with the fNIRS topographies, it can be concluded that the limited depth sensitivity of the fNIRS measurements did not introduce a substantial bias to the present results. The current study thus demonstrates that fNIRS is a suitable method for investigating auditory cortex activity, both in terms of measurement depth and spatial resolution.

## V. CONCLUSIONS

The current results have shown that speech-like acoustic input with slowly changing prosodic features primarily evokes activity in the right auditory cortex of normal-hearing adult listeners, in line with previous studies. This finding was evident in the fNIRS HbR and EEG data on group level as well as the subgroup comparison of the fNIRS HbO data. The agreement of the results of both methods demonstrates that fNIRS is a suitable tool for investigating auditory perception.

On group level, however, the fNIRS HbO results deviated markedly from this pattern as they were dominated by a prominent anterior-to-posterior blood flow gradient that masked the neural activity in the auditory cortex, resulting in negative responses in auditory areas. As this gradient was not observed in the non-cortical superficial tissue layers and the time course of these negative responses was slower than that of a typical hemodynamic response elicited by neural activity, the gradient appears to be caused by large-scale cortical blood flow changes, so-called ‘blood stealing’.

To further investigate this effect, the subjects were divided into subgroups based on their HbO responses in the right auditory cortex. Unexpectedly, the EEG data suggested that positive HbO responses in this region were accompanied by smaller auditory ERPs and reduced attention. The analyses of the concurrently recorded EEG data hence revealed that larger fNIRS HbO responses are not necessarily a positive finding when investigating auditory perception, contrary to what might be expected. More generally, the fact that the different EEG analyses have helped to avoid this potential fallacy shows that the benefits of combining fNIRS and EEG go far beyond the common notion of joining the good spatial resolution of blood-based measures with the good temporal resolution of electrophysiological data.

## ACKNOWLEDGEMENTS

We are grateful to the Dietmar Hopp Stiftung (Grant No. 2301 1239) for supporting our research. We would like to thank Stuart Rosen for providing the stimulus materials, and Bettina Sorger and Alexander Gutschalk for helpful advice regarding data analysis and the interpretation of the results.

